# Bortezomib inhibits lung fibrosis and fibroblast activation without proteasome inhibition

**DOI:** 10.1101/2021.02.26.433086

**Authors:** Loka Raghu Kumar Penke, Jennifer Speth, Scott Wettlaufer, Christina Draijer, Marc Peters-Golden

## Abstract

The FDA-approved proteasomal inhibitor bortezomib (BTZ) has attracted interest for its potential anti-fibrotic actions. However, neither its *in vivo* efficacy in lung fibrosis nor its dependence on proteasome inhibition has been conclusively defined. Herein, we identify that therapeutic administration of BTZ in a mouse model of pulmonary fibrosis diminished the severity of fibrosis without reducing proteasome activity in the lung. Under conditions designed to mimic this lack of proteasome inhibition *in vitro*, it reduced fibroblast proliferation, differentiation into myofibroblasts, and collagen synthesis. It promoted de-differentiation of myofibroblasts and overcame their characteristic resistance to apoptosis. Mechanistically, BTZ inhibited kinases important for fibroblast activation while inducing expression of dual-specificity phosphatase 1 or DUSP1, and knockdown of DUSP1 abolished its anti-fibrotic actions in fibroblasts. Our findings identify a novel proteasome-independent mechanism of anti-fibrotic actions for BTZ and support its therapeutic repurposing for pulmonary fibrosis.

## Introduction

Fibrotic diseases are responsible for extensive morbidity and mortality worldwide. Idiopathic pulmonary fibrosis (IPF), the most common fibrotic disease of the lung, often progresses to respiratory failure and death within 2-3 years from the time of diagnosis. Two FDA-approved drugs, nintedanib and pirfenidone, delay disease progression, but neither reverses established IPF nor improves patient survival (1, 2). These realities highlight the unmet medical need for the development of new treatments for IPF. Fibroblasts (Fibs) are the principal “effector” cells of fibrotic disorders by virtue of their capacity to proliferate and to differentiate into myofibroblasts (MyoFibs). MyoFibs are marked by their expression and organization of α-smooth muscle actin (α-SMA) into contractile stress fibers, and their pathogenic importance derives from their capacities to elaborate excessive amounts of extracellular matrix proteins such as collagen that comprise scars and to exhibit a high degree of resistance to apoptosis (3).

MyoFib differentiation is characterized by reprogramming of the global transcriptome and proteome. Concomitant with an increase in new protein synthesis is the need to degrade unwanted proteins via the ubiquitin-proteasome system. Indeed, increased activity of this proteolytic complex has been reported in fibrotic lung (4–6) and other organs (7), and in Fibs treated with pro-fibrotic mediators (6, 8, 9). At the molecular level, increased expression of ubiquitin ligases as well as proteasome subunits and their activators has been reported in fibrotic tissue and Fibs, including those from IPF patients (10–12). These findings provide a potential therapeutic rationale for proteasome inhibition in fibrosis.

Bortezomib (BTZ) is a reversible inhibitor of the chymotrypsin-like activity of the 20S core proteasome. The first FDA-approved proteasome inhibitor, it is indicated for the treatment of multiple myeloma. BTZ and other proteasome inhibitors have been evaluated in experimental models of fibrosis in a variety of tissues (13–18). While BTZ was found to ameliorate fibrosis in other organs, data on its efficacy in models of pulmonary fibrosis have been limited and conflicting (16, 17). Moreover, whether its potential anti-fibrotic actions actually depend on its proteasomal inhibitory capacity has never been explicitly determined.

In this study, we demonstrate that BTZ exerts robust and potent anti-fibrotic actions both *in vitro* and *in vivo* in the absence of proteasome inhibition, and identify induction of dual-specificity phosphatase 1 (DUSP1) as a potential new mechanism for these actions.

## Methods

### Cell culture

CCL-210 (CCD-19Lu) and MRC5 (CCL171) adult human lung Fib lines were purchased from ATCC. For selected studies, we employed Fibs grown from biopsy specimens of patients at the University of Michigan determined to have either IPF or non-fibrotic lung under an IRB-approved protocol, as reported previously (5). Fibs were cultured in low glucose Dulbecco’s modified Eagle’s medium (DMEM) and supplemented with 10% fetal bovine serum (Hyclone) and 100 units/ml of both penicillin and streptomycin (Invitrogen), hereafter referred as complete medium.

### Reagents

Recombinant human transforming growth factor-β (TGF-β) and fibroblast growth factor-2 FGF-2 were purchased from R&D Systems. BTZ, bleomycin and human activating FAS (anti-FAS) antibody were purchased from Millipore Sigma. PGE_2_, forskolin and DRB were purchased from Cayman Chemicals. Unless otherwise specified, the final concentrations of modulatory agents used for cell treatment were: TGF-β, 2 ng/ml; FGF-2, 50 ng/ml; BTZ, 10 nM; PGE_2_, 500 nM; forskolin, 10 μM; and DRB, 25 μM.

### RNA isolation and quantitative real-time PCR

Cells were lysed in 700 μl TRIzol reagent (Thermo Scientific) and total RNA was extracted using the RNeasy Mini Kit (Qiagen). RNA was estimated using Nanodrop and converted to cDNA using the high capacity cDNA reverse transcription kit (Applied Biosystems). Gene expression was quantified using Fast SYBR green master mix (Applied Biosystems) on a StepOne Real-time PCR system (Applied Biosystems). Expression studies for human *FOXM1*, *CCNB1*, *PLK1*, *CCND1*, *Survivin*, *α-SMA*, *COL1α2*, *APAF1*, *BID*, *FASR*, *COX2, DUSP1* and mouse *Col1α1*, *Ctgf* and *TGF-β1* were performed using specific primers listed in **Supplementary Table 1 and 2**. Relative quantification of gene expression was determined using the ΔCT method, and GAPDH and β-actin were used as reference genes for human and mouse samples, respectively.

### Western blot

Samples were lysed in RIPA buffer (Cell Signaling) supplemented with protease inhibitors (Roche Diagnostics) and phosphatase inhibitor cocktail (EMD Biosciences). Sources of antibodies were as follows: ubiquitin and *α*-SMA, Abcam; FAS and collagen 1, Thermo Scientific; MKP1 and FOXM1, Millipore; and Cyc B1, P38, phosphoP38, Akt, pAkt, PARP, and GAPDH-HRP conjugate, Cell Signaling Technologies. All antibodies were used at a dilution of 1:1,000 except GAPDH-HRP which was used at a dilution of 1:3000. Protein quantification was performed using ImageJ software.

### Cell viability assay

Cytotoxicity of BTZ in CCL-210 cells was assessed using a luciferase-coupled ATP quantitation assay (CellTiter-Glo from Promega). Briefly, cells were seeded in a 96-well plate at a density of 5 × 10^3^ cells/well in complete medium overnight. Cells were then treated with different doses of BTZ (4 - 512 nM). After 72 h, culture medium was removed, washed with PBS, and 100 μL of CellTiter-Glo reagent was added to each well and incubated for 30 min at room temperature. Luminescence intensity was measured at 450 nm on a Tecan infinity 200 PRO.

### Fib proliferation

Proliferation studies were performed using the CyQUANT NF Cell Proliferation Assay Kit (Life Technologies). Briefly, Fibs were plated at 5 × 10^3^ cells/well in a 96-well plate, allowed to adhere overnight (16 h) and then shifted to serum-free medium for 24 h. Cells were then treated with BTZ (10 nM) for 30 min and after a change of medium, stimulated with FGF-2 at 50 ng/ml in serum-free DMEM for 72 h at 37°C. After removing the medium, 100 μl of 1X Hank’s Balanced Salt Solution containing CyQuant NF dye was added to each well and incubated at 37°C for 45 min. Fluorescence was measured using a fluorescence microplate reader with excitation at 485 nm and emission at 530 nm.

### Fib differentiation

Fibs were incubated with TGF-β at 2 ng/ml for 48 h to differentiate them into myofibroblasts. Differentiation was confirmed by determining expression of *α*-SMA and COL1*α*2 (by qPCR and Western blot).

### Apoptosis

Apoptosis in Fibs, MyoFibs, and IPF Fibs was evaluated at baseline and after stimulating with anti-FAS antibody. Apoptosis was determined by measuring i) cell-surface expression of phosphatidylserine using annexin V–FITC staining as determined by flow cytometry, ii) caspase 3/7 activity, and iii) expression of pro-survival and pro- apoptotic genes and of the death receptor FAS.

### cAMP ELISA

cAMP levels in cell lysates were determined using an ELISA kit (Enzo Life Sciences) according to the manufacturer’s protocol. Briefly, cells were treated with 10 nM BTZ as detailed under the Results section and lysed by incubating with 0.1M HCl for 20 min. Lysates were then centrifuged at 1500 x *g* for 10 min, and cAMP levels measured in the supernatant.

### PGE_2_ ELISA

Supernatants from CCL-210 cells treated with or without 10 nM BTZ were harvested at various timepoints. Samples were then centrifuged at 1500 x *g* for 10 min and the levels of PGE_2_ were quantified from the cell-free culture supernatants by a PGE_2_ ELISA kit (Enzo Life Sciences) according to the manufacturer’s instructions.

### Bleomycin model of pulmonary fibrosis

Studies were approved by the University of Michigan Committee on Use and Care of Animals. 6-8 week old female C57BL/6 mice (Charles River Laboratories) received a single oropharyngeal dose of 1.5 units/kg of bleomycin or an equal volume of saline. BTZ was administered i.p. at 0.1 or 0.25 mg/kg beginning at day 9 post-bleomycin and every 3 days thereafter. Mice were sacrificed on day 21 and lungs were harvested to study fibrotic end points. The left lung was analyzed for hydroxyproline content as described previously, while the right lung lobes were assessed for the expression of fibrotic marker genes (*Col1a1, Ctgf and TGF-β1*), Masson’s trichrome staining, and 20S proteasome activity.

### 20S proteasome activity assay

CCL-210 cells were seeded in 60 mm petri dishes at a density of 5 × 10^5^ cells/well in complete medium. After 24 h, cells were treated with different doses of BTZ or an equal volume of DMSO, as depicted in the experimental layout shown in Figure 2A. Samples were harvested at designated time points, washed with 20S Proteasome Assay buffer and lysed with 20S Proteasome Lysis Buffer (both from Cayman Chemicals) and frozen at -80°C until assay. Samples were thawed on ice and incubated at room temperature for 30 min on an orbital shaker. To determine the proteasome activity in mouse lungs, lung tissue was homogenized, and samples frozen at -80°C until assay.

**Figure 1.**
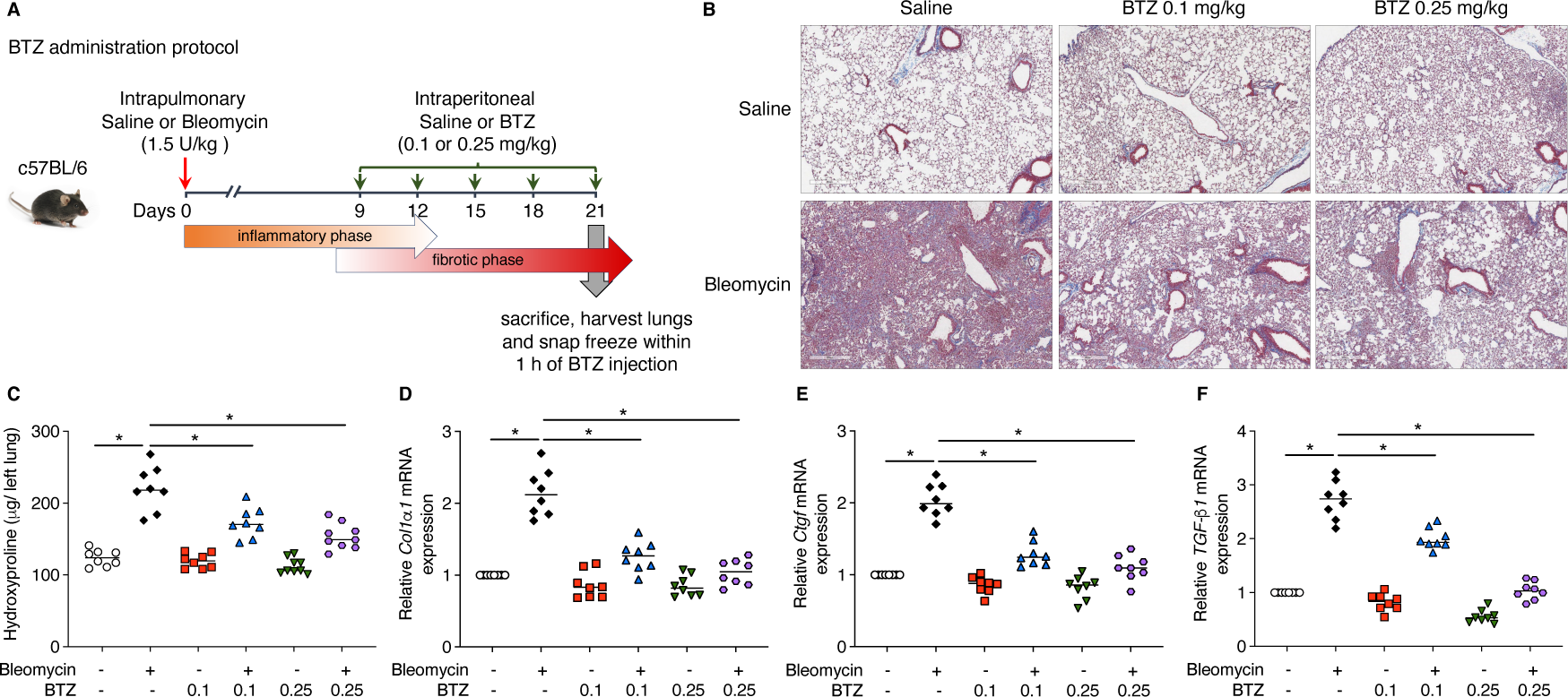
BTZ administration improves bleomycin-induced fibrosis in mice. (**A**) Scheme illustrating the timelines for *in vivo* administration of bleomycin and BTZ, for determination of experimental endpoints at day 21, and for the pertinent phases of the pulmonary response in the bleomycin model of pulmonary fibrosis. (**B**) Digital images of Masson’s trichrome staining for collagen deposition (blue) at day 21 in mice treated ± bleomycin and BTZ. Original magnification, ×200. Scale bars: 500 μm. (**C-F**) Effect of BTZ treatment in mice treated ± bleomycin as reflected by changes in lung hydroxyproline content (**C**) and the mRNA expression of fibrotic markers *Col1*α*1*, *Ctgf*, and *Tgf-β1* (**D-F**); in (**C-F**), each symbol represents an individual mouse and horizontal lines represent mean values. Values in each group represent results from two pooled independent experiments with a total of 8-9 mice per group. \**P* < 0.05; two-way ANOVA.

**Figure 2.**
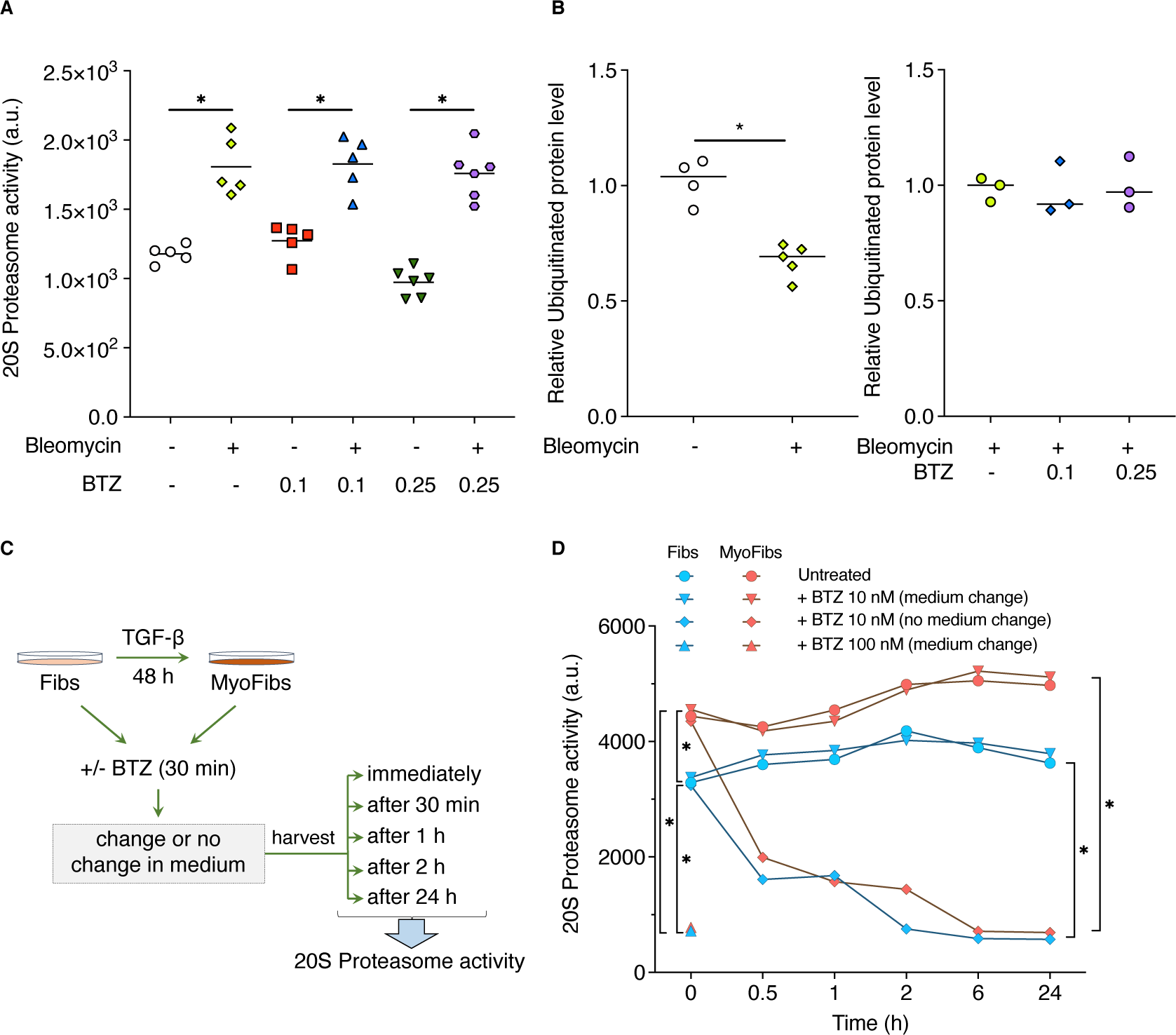
Influence of BTZ on proteasome activity *in vivo* and *in vitro*. (**A**) Lung tissues from *in vivo* groups in Figure 1 were assessed for proteasome activity via 20S proteasome activity assay 1 h after harvest. (**B**) Densitometric analysis of ubiquitinated protein bands in lung tissue harvested at day 21 from mice treated with saline or bleomycin (left) or bleomycin alone or bleomycin followed by treatment with BTZ 0.1 mg/kg or 0.25 mg/kg (right); total density of the lane for each mouse is expressed relative to the density of the GAPDH band for that lane. (**C**) Design of experiments to assess effect of BTZ dose and incubation protocol on proteasome activity in Fibs and in MyoFibs. (**D**) Fibs and MyoFibs were treated with BTZ at 10 or 100 nM and medium was either changed at 30 min or not changed, and 20S proteasome activity was assessed immediately or at 30 min, 1 h, 2 h, 6 h, or 24 h later. In **(A)**, each symbol represents an individual mouse and horizontal lines represent mean values. Each symbol in (**B**) represents individual mice with mean values. Values in (**D**) represent mean values (± S.E.) from 3 independent experiments. For (**A**) and (**D**), \**P* < 0.05; two-way ANOVA and for (**B**) **P < 0.01; Student’s t test, unpaired.

Samples were thawed on ice prior to assay. Sample lysates generated from *in vitro* Fib cultures or lung tissue harvested following *in vivo* experiments were centrifuged (1500 x g for 10 min at 4°C) and 90 µl of clear supernatant from each sample was transferred into each well of a 96-well black plate and proteasome activity was measured using Suc-LLVY-AMC substrate (Millipore) according to the manufacturer’s instructions.

### CRISPR/Cas9-mediated knockdown

MRC5 lung Fibs were used for guide RNA-based DUSP1 knockdown. Guide RNA against DUSP1 (sgDUSP1) or non-targeting control (sgCont) and Cas9 2NLS nuclease were purchased from Synthego. We mixed sgRNAs and Cas9 at a ratio of 1.3:1 in Opti-MEM (Invitrogen) according to the manufacturer’s instructions to prepare ribonucleoprotein complexes. We then added Lipofectamine CRISPRMAX in Opti-MEM to generate a transfection mix, incubated for 15 min at room temperature and then added to the cells. Cells were cultured for 48 h to achieve *DUSP1* KD.

### Statistics

Unless specified otherwise, all data were from a minimum of 3 independent experiments. Data were reported as mean ± s.e.m. Group differences were compared using the unpaired two-sided Student’s *t*-test or two-way ANNOVA with post hoc Tukey’s correction for multiple comparisons, as appropriate. A *P* value < 0.05 was considered statistically significant.

## Results

### BTZ attenuates bleomycin-induced lung fibrosis in mice

BTZ was administered at 0.1 and 0.25 mg/kg i.p. every third day beginning at day 9 post-bleomycin, and lungs were harvested on day 21 and samples were snap-frozen in liquid nitrogen within 1 h after the last BTZ injection (**Figure 1A**) for biochemical studies. As expected, bleomycin challenge resulted in collapse of alveoli with marked deposition of interstitial collagen as revealed by trichrome staining demonstrated in both low-magnification images (**Supplementary Figure 1)** and high-magnification images (**Figure 1B**). The increased collagen deposition was confirmed biochemically by increased levels of hydroxyproline (**Figure 1C**). Expression of fibrotic marker genes *Ctgf*, *Col1α1*, and *Tgf-β1* (**Figures 1D-F**) was also increased. In the absence of bleomycin, neither dose of BTZ alone had any impact on any of the endpoints examined. However, both doses of BTZ significantly and substantially reduced all of the fibrotic endpoints, with the dose of 0.1 mg/kg achieving a near-maximal effect (**Supplemental Figure 1** and **Figures 1B-F**).

### The anti-fibrotic actions of BTZ are independent of proteasome inhibition

To evaluate proteasome activity in the lung tissues harvested at day 21, we utilized snap-frozen lung tissue prepared within 1 h after the last BTZ or saline injection. We chose this 1 h interval based on pharmacokinetic data demonstrating the transient nature of enzymatic inhibition by BTZ, with ∼ 75% inhibition observed at 1 h after dosing (19–21). Prior studies revealed that BTZ selectively inhibits the chymotrypsin-like peptidase activity of the β5 subunit of the 20S proteasome catalytic core (22–24). Therefore, we chose to evaluate lung tissue proteasome activity using a commercially available 20S proteasome assay kit sensitive to chymotrypsin activity. Consistent with prior findings (5, 6), fibrotic lungs following bleomycin challenge exhibited increased proteasome activity as compared to saline-challenged controls. However, treatment with BTZ at either dose had no effect on proteasome activity in the lungs of either bleomycin- or saline-challenged mice (**Figure 2A**). Proteasome inhibition is anticipated to lead to increased accumulation of ubiquitinated proteins, but no such increase was observed by immunoblotting with an anti-ubiquitin antibody in lungs from mice treated with BTZ at either dose in the lungs of bleomycin-challenged mice (**Figure 2B**, right and **Supplementary Figure 2B**). However, the significant reduction in global ubiquitinated protein content in lungs from bleomycin-treated mice as compared to control mice (**Figure 2B**, left) corroborates the increased proteasomal activity observed in the former group (**Figure 2A** and **Supplementary Figure 2A**). Thus, our data disassociate the anti-fibrotic actions of BTZ seen in **Figure 1** from its inhibition of proteasome function.

A number of reports of BTZ actions in Fibs employed doses of 10 nM-1 μM along with continuous exposure for prolonged time intervals of 24-48 h (16, 17, 25). To define conditions in Fibs (or MyoFibs established by 48 h pretreatment with TGF-β) in which BTZ does and does not inhibit the proteasome, we examined a range of doses and also took advantage of the reversibility of its known proteasome inhibitory actions. This entailed measuring 20S proteasome activity at various time points in lysates of Fibs or MyoFibs treated either continuously or for only 30 min with BTZ before replacing the culture medium, as illustrated in **Figure 2C**. Consistent with an increased proteasome activity observed in lung from bleomycin-treated mice (**Figure 2A**), MyoFibs elicited *in vitro* by TGF-β treatment likewise showed a significant increase in proteasome activity as compared to that of Fibs (**Figure 2D**). We observed substantial inhibition of proteasome activity at 30 min with a BTZ concentration of 100 nM, and a time-dependent inhibition with 10 nM BTZ in cells exposed without a medium change. In contrast, when incubated with cells at 10 nM for only 30 min before washing and medium change, BTZ exerted no effect on proteasome activity at time points ranging from 30 min to 24 h. Notably, the effects of BTZ on proteasome activity were qualitatively the same in Fibs and MyoFibs. In separate studies we evaluated the effects of a 30-min treatment with BTZ on Fib cytotoxicity and found that BTZ concentrations <64 nM had no effect on Fib viability whereas those ≥64 nM reduced viability in a concentration-dependent manner (**Supplementary Figure 3**). We therefore chose a 30-min exposure with 10 nM BTZ and subsequent medium change to obtain conditions that neither inhibited the proteasome nor elicited cytotoxicity – thereby mimicking the BTZ effects observed in lung tissue *in vivo* – in order to assess various Fib activation properties *in vitro*.

### BTZ inhibits FGF-2-induced Fib proliferation

We used these conditions to assess the effects of BTZ on cell proliferation and expression of relevant cell cycle-related transcripts and proteins in Fibs treated ± the mitogen FGF-2 (**Figure 3A**). Consistent with our prior studies (5), stimulation with FGF-2 increased cell proliferation (**Figure 3B**), as well as mRNA and protein expression of the key proliferation-regulatory transcription factor FOXM1 and the FOXM1-dependent cell cycle gene *CYCB1*. All of these parameters were significantly inhibited by pretreatment with BTZ (**Figure 3C-3D**).

**Figure 3.**
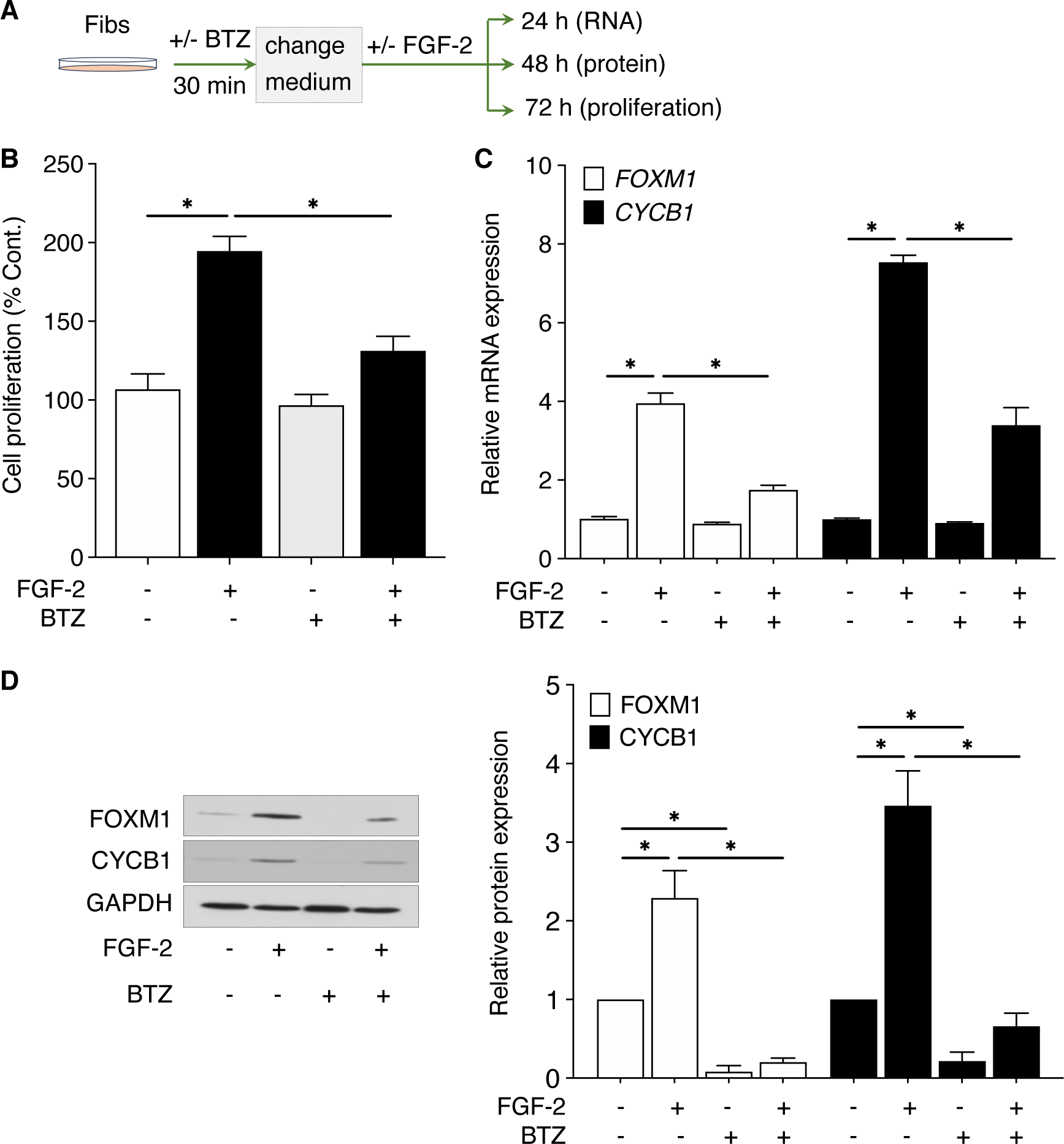
BTZ inhibits FGF-2-induced Fib proliferation. (**A**) Design of experiments to evaluate BTZ effects on FGF-2-induced Fib proliferation endpoints depicted in (**B**-**D)**. (**B**) Cells were pretreated +/-BTZ (10 nM) for 30 min, after which the medium was replaced, and they were stimulated +/-FGF-2. Cells were harvested and proliferation was quantified at 72 h. Control value represents fluorescence of DMSO-treated Fibs. (**C**-**D**) Cells were pretreated +/-BTZ (10 nM) for 30 min, after which the medium was replaced, and they were stimulated +/-FGF-2. Cells were harvested and assessed for the expression of proliferation-associated genes *FOXM1* and *CYCB1* at 24 h (**C**), and for the expression of FOXM1 and CYCB1 proteins by Western blot at 48 h (**D**); left panel, representative blot; right panel, mean densitometric analysis of blots from 3 experiments. All data represent mean values (± S.E.) from 3 independent experiments. *P < 0.05, two-way ANOVA.

### BTZ prevents TGF-β-induced Fib differentiation

We assessed whether BTZ could prevent differentiation of Fibs into MyoFibs elicited by treatment with TGF-β (**Figure 4A**). Stimulation with TGF-β resulted in increased mRNA and protein expression of α-SMA and Col1α2 (**Figure 4B-4C**), characteristic phenotypic features of MyoFibs. Pretreatment with BTZ significantly and markedly reduced both baseline as well as TGF-β-stimulated expression of α-SMA and Col1α2 at both mRNA and protein levels (**Figure 4B-4C**).

**Figure 4.**
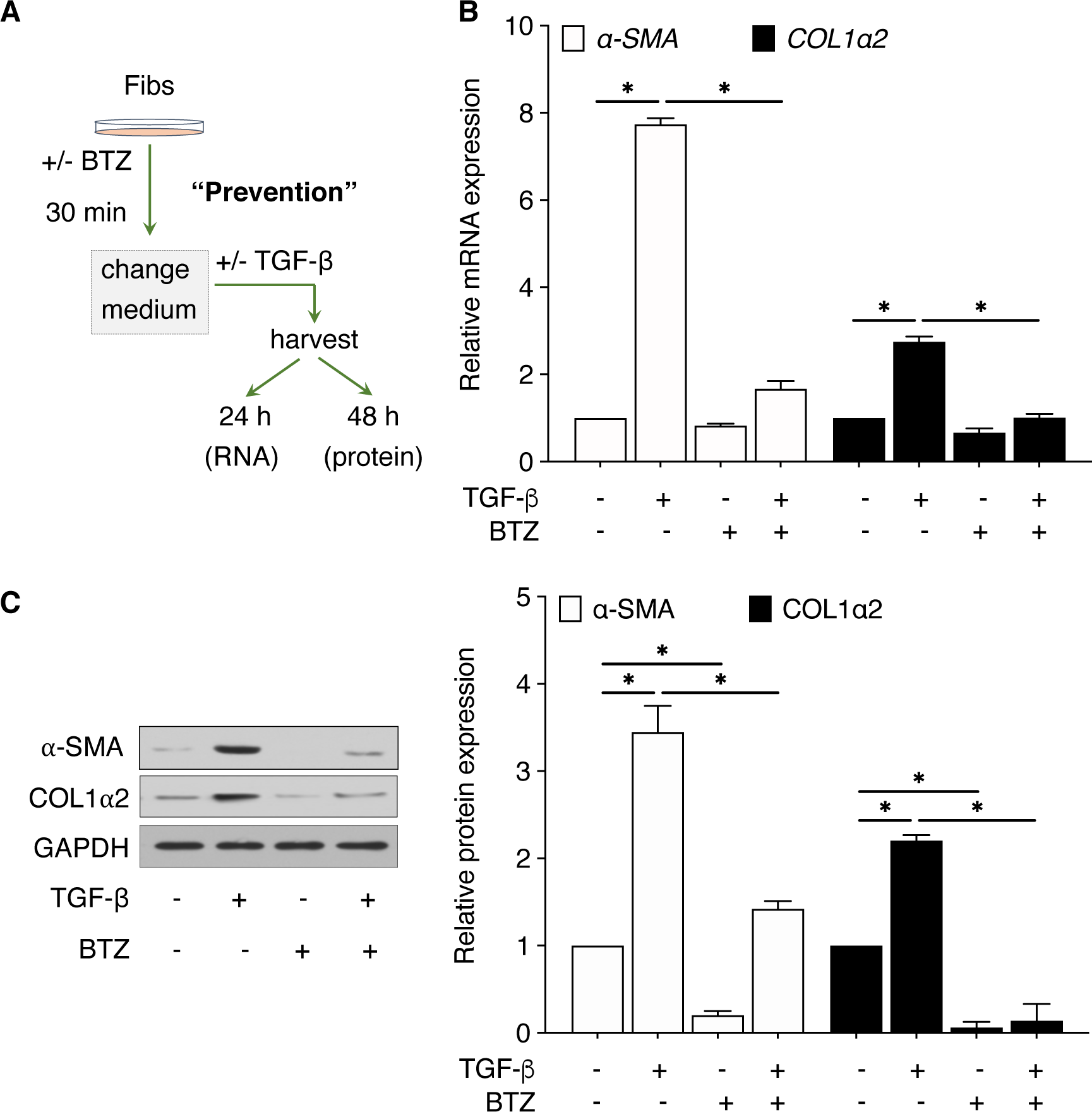
BTZ prevents TGF-β-induced Fib differentiation. (**A**) Design of experiments to assess the capacity of BTZ to attenuate Fib differentiation using a “prevention protocol”. (**B-C)** Fibs were treated +/- BTZ (10 nM) for 30 min, after which the medium was replaced, and cells stimulated +/- TGF-*β*. Cells were harvested and assessed for the expression of differentiation-associated genes *α-SMA* and *COL1α2* mRNA by qPCR at 24 h (**B**) and their proteins by Western blot at 48 h (**C**). In (**C**), the left panel depicts a representative blot, and the right panel provides densitometric analysis of blots from 3 experiments. All data represent mean values (± S.E.) from 3 independent experiments. *P < 0.05, 2-way ANOVA.

### BTZ promotes de-differentiation of elicited MyoFibs and IPF Fibs

We investigated the capacity of BTZ to reduce α-SMA and Col1α2 expression in MyoFibs already established by 48 h pretreatment with TGF-β – a phenomenon termed “de-differentiation” (**Figure 5A**). BTZ significantly decreased expression of α-SMA and Col1α2 at the mRNA and protein levels (**Figure 5B** and **5C**). The basal expression levels of α-SMA are significantly higher in Fibs from patients with IPF than in non-fibrotic control Fibs (5), reflecting a baseline degree of MyoFib differentiation. BTZ treatment in IPF lines is illustrated in **Figure 6A**. Treatment of IPF Fib lines with BTZ similarly reduced baseline expression of α-SMA and Col1α2 mRNA (**Figure 6B**) and protein (**Figure 6C**), reflecting active de-differentiation.

**Figure 5.**
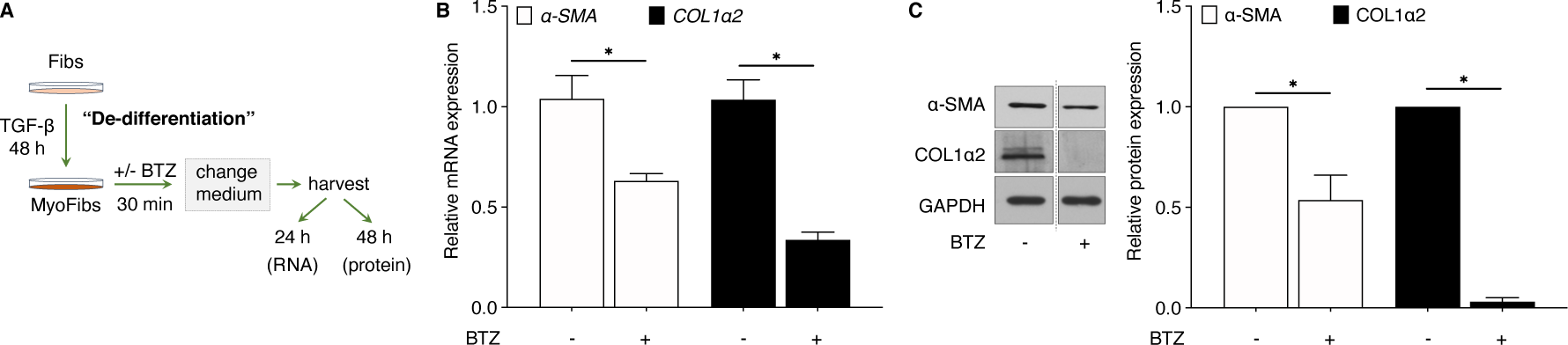
BTZ de-differentiates established TGF-β-elicited MyoFibs. (**A**) Design of experiments to assess the capacity of BTZ to reduce MyoFib phenotype in a “de-differentiation protocol”. (**B**-**C)** Fibs treated for 48 h with TGF-*β* to elicit differentiation into MyoFibs were then treated +/- BTZ (10 nM) for 30 min, after which the medium was changed. Cells were harvested and assessed for the expression of *α- SMA* and *COL1α2* mRNA by qPCR at 24 h (**B**) and protein by Western blot at 48 h (**C**). In (**C**), the left panel presents a representative blot and the right panel provides mean densitometric values (±S.E.) of Western blots from 3 independent experiments. For (**B**) and (**C**), GAPDH mRNA and protein were used to normalize α-SMA and COL1α2 expression by qPCR and Western blot, respectively. All data represent mean values (± S.E.) from 3 independent experiments. *P < 0.05, two-way ANOVA.

**Figure 6.**
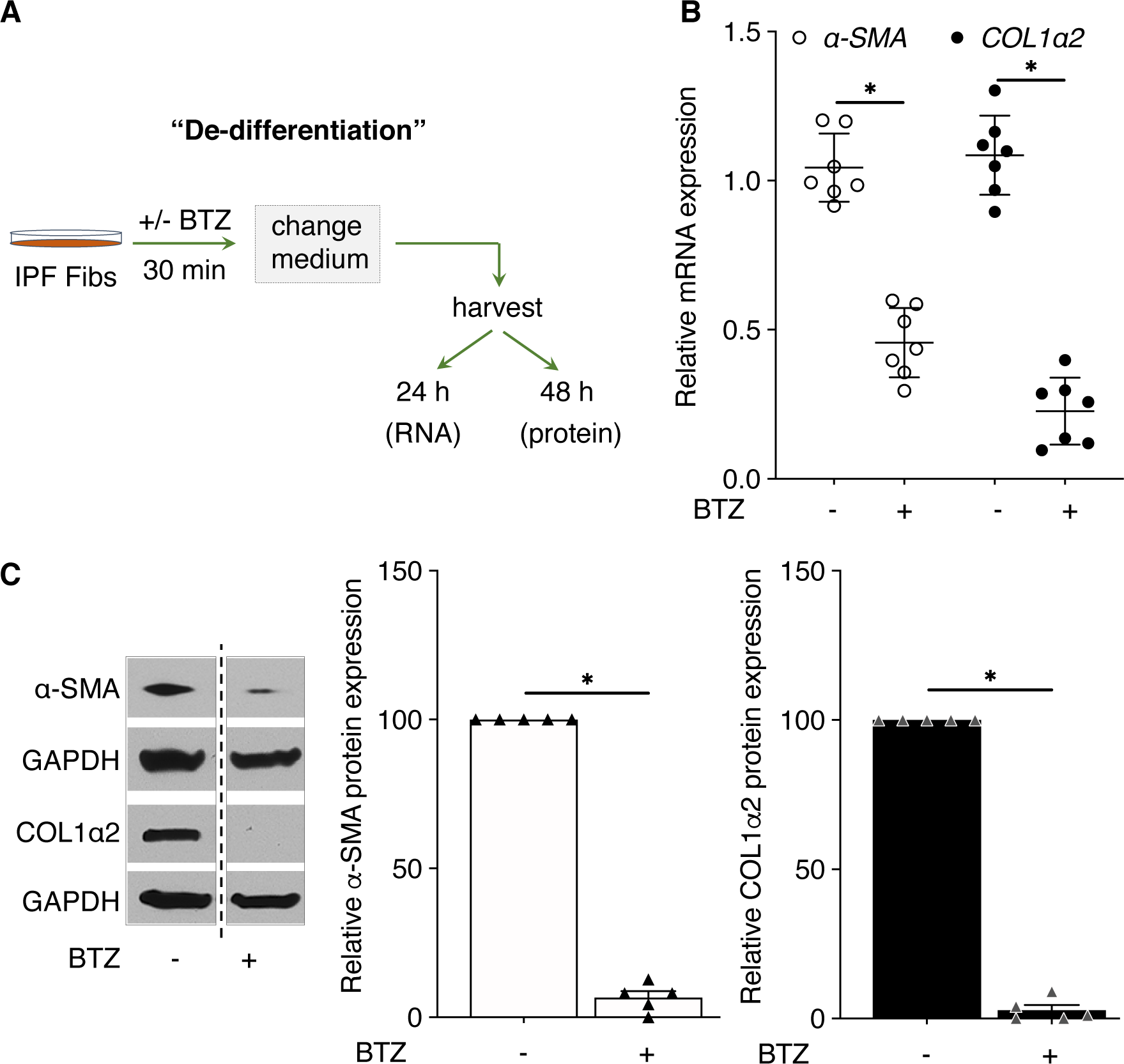
BTZ de-differentiates IPF Fibs. (**A**) Design of experiments to assess BTZ effects on IPF Fibs using a combination “de- differentiation” and “prevention” protocol. (**B**-**C**) IPF Fibs were treated +/- BTZ (10 nM) for 30 min, after which the medium was replaced, and they were stimulated +/- TGF-*β*. Cells were harvested and analyzed for the expression of *α-SMA* and *COL1α2* mRNA by qPCR at 24 h (**B**) and protein by Western blot at 48 h (**C**). For (**B**) and (**C**), GAPDH mRNA and protein were used to normalize α-SMA and COL1α2 expression by qPCR and Western blot, respectively. In (**C**), the left panel presents a representative blot and the middle and right panels depict mean densitometric analysis of α-SMA and COL1α2 from Western blots from 3 experiments; the dashed line in the left panel indicates that the lanes were from the same blot but non-contiguous. All data represent mean values (± S.E.) from 3 independent experiments. *P < 0.05, two-way ANOVA.

### BTZ sensitizes MyoFibs and IPF Fibs to FAS-mediated apoptosis

Increased resistance to apoptosis, as compared to that of Fibs, is a pathophysiologically relevant hallmark of MyoFibs (3, 26). We sought to determine if the de-differentiation of MyoFibs achieved by BTZ treatment sensitized them to apoptosis. **Figure 7A** demonstrates the expected susceptibility of Fibs to apoptosis, assessed by measuring Annexin V binding in response to a 14-h incubation with an antibody (anti-FAS) that activates the death receptor FAS. By contrast, TGF-*β*-elicited MyoFibs were highly resistant to FAS-mediated apoptosis (**Figure 7C**). However, treatment of these MyoFibs with BTZ using the de-differentiation protocol depicted in **Figure 5A** prior to addition of anti-FAS (**Figure 7B**) rendered them highly susceptible to FAS-mediated apoptosis (**Figure 7C**). The ability of BTZ to promote FAS-mediated apoptosis in MyoFibs was also confirmed independently by caspase 3/7 activity (**Figure 7D**). Mechanistically, this restoration of apoptosis susceptibility by BTZ was associated with a reduction in the expression of the pro-survival *BIRC5*, an increase in pro-apoptotic genes *APAF1* and *BID* (**Figure 7E**), and an increase in the mRNA and protein levels of FAS itself (**Figure 7F**). These data show that de-differentiation of MyoFibs elicited by BTZ markedly enhances their susceptibility to apoptosis via multiple molecular mechanisms. Next, as illustrated in **Figure 7G**, we determined caspase 3/7 activity on IPF Fibs treated with BTZ to determine its effect on apoptosis. As it did for TGF-*β*-elicited MyoFibs, BTZ also sensitized otherwise resistant IPF Fibs to apoptosis elicited by anti-FAS (**Figure 7H**). Collectively, our findings demonstrate that BTZ strongly sensitizes Fibs and MyoFibs to FAS-mediated apoptosis.

**Figure 7.**
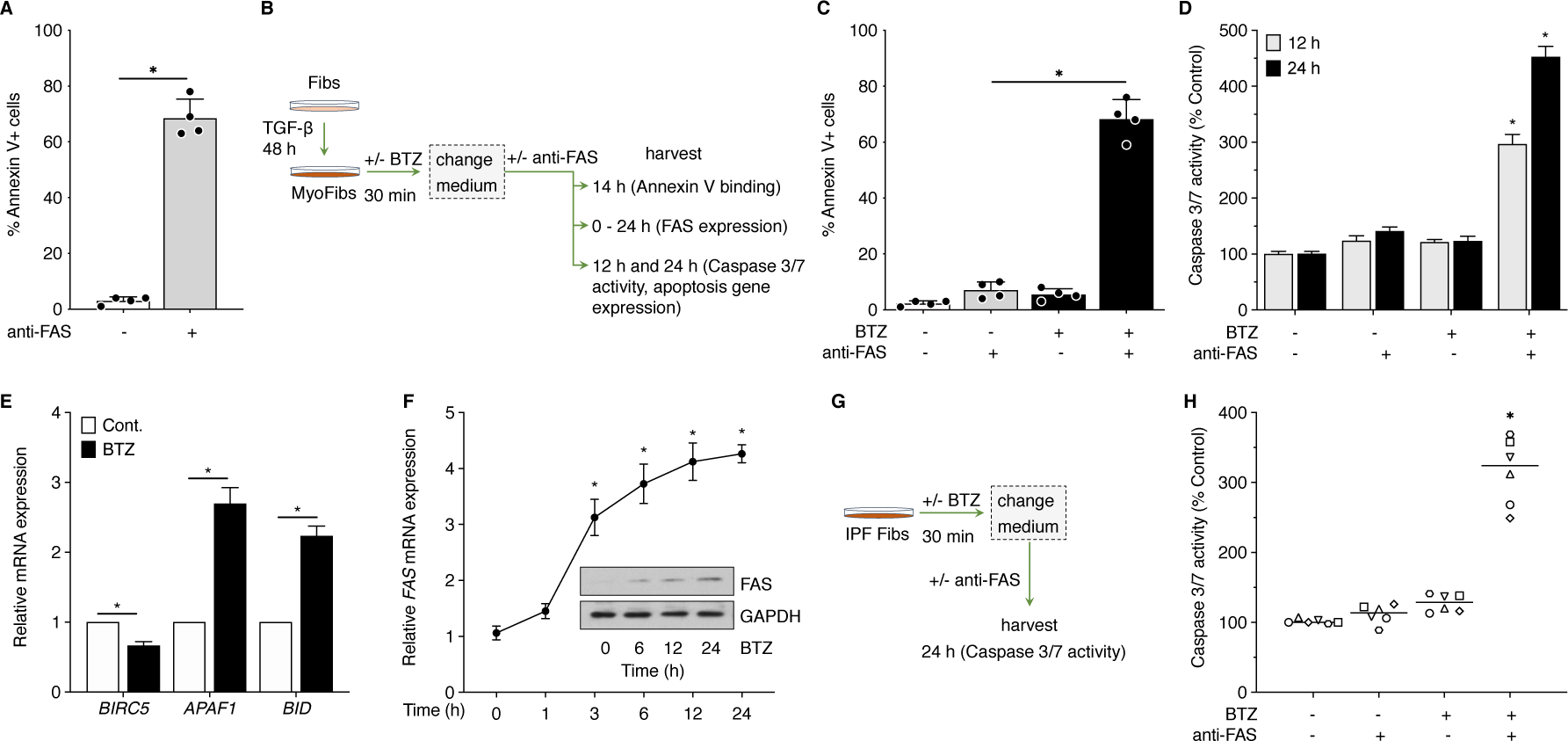
BTZ sensitizes established MyoFibs to FAS-mediated apoptosis. (**A**) Fibs were treated with activating anti-FAS antibody at 100 ng/ml for 14 h. Cells were harvested, phosphatidylserine on early apoptotic cells was detected by Pacific Blue- Annexin-V staining, and the percentage of apoptotic cells was quantified by flow cytometry. (**B**) Design of experiments to assess BTZ effect on TGF-*β*-elicited MyoFib apoptosis. (**C**-**F**) TGF-*β*-elicited MyoFibs were treated +/- BTZ (10 nM) for 30 min, after which the medium was changed, and they were then stimulated +/- anti-FAS antibody as in (**A**). Cells were harvested and apoptotic cells quantified either by flow cytometric analysis of phosphatidylserine staining at 14 h (**C**) or by caspase 3/7 activity assay (**D**). (**E**) Effect of BTZ on expression of the pro- and anti-apoptotic genes *BIRC5, APAF1, and BID* measured by qPCR at 24 h. (**F**) Effect of BTZ on expression of FAS mRNA by qPCR and FAS protein analysis by Western blot at the time points indicated. (**G**) Design of experiments to assess BTZ effect on IPF Fib apoptosis sensitivity. (**H**) IPF Fibs (n = 6) were treated +/- BTZ (10 nM) for 30 min, after which the medium was changed, and stimulated +/- anti-FAS antibody. Cells were harvested at 24 h and apoptosis was quantified by caspase 3/7 activity. In (**E-F**), GAPDH mRNA and protein were used to normalize apoptosis genes or FAS expression by qPCR and Western blot, respectively. All data represent mean values (± S.E.) from 3 independent experiments. *P < 0.05, two-way ANOVA.

### Inhibition of Fib activation by BTZ is independent of prostaglandin production or signaling

The pleiotropic inhibitory actions of BTZ displayed in **Figures 3-7** – including prevention of Fib proliferation and MyoFib differentiation, de-differentiation of established MyoFibs, and sensitization to apoptosis – are strikingly reminiscent of effects previously reported for the endogenous lipid mediator prostaglandin E_2_ (PGE_2_) by our laboratory and others (27–30). As shown in **Supplementary Figure 4A**, we considered the possibility that BTZ might mediate these diverse actions by promoting PGE_2_ synthesis or its signaling via the intracellular second messenger cyclic AMP (cAMP). First, we examined the kinetics of mRNA expression of cyclooxygenase-2 (COX-2), a pivotal enzyme in PGE_2_ biosynthesis, in Fibs treated ± BTZ. BTZ had no effect on COX-2 gene expression, whereas PGE_2_, which is known to induce COX-2 expression in a positive feedback loop, elicited robust induction of COX-2 (**Supplementary Figure 4B**). Next, we utilized ELISA to directly measure the amounts of PGE_2_ produced and secreted into the conditioned medium over a 24 h incubation ± BTZ. Again, BTZ failed to meaningfully increase Fib PGE_2_ generation, in contrast to the direct adenylyl cyclase activator forskolin, which did so to a marked extent (**Supplementary Figure 4C**). The Fib-inhibitory actions of PGE_2_ are mediated by increases in intracellular cAMP (29, 31), so we next considered the possibility that BTZ might amplify intracellular cAMP levels independent of increases in PGE_2_ generation; however, as shown in **Supplemental Figure 4D**, while forskolin increased cAMP levels as expected, BTZ had no such effect. Together, these data argue that increased PGE_2_ synthesis or cAMP signaling is unlikely to underlie the Fib- inhibitory actions of BTZ.

### BTZ inhibits key kinases activated by TGF-β and FGF-2

We sought to define a proteasome-independent mechanism to explain the multidimensional deactivation of Fib functions by BTZ. Phosphorylation and concomitant activation of P38 and AKT, respectively, have been implicated in TGF-β-mediated differentiation (29) and FGF-2-mediated proliferation (5) of lung Fibs. We therefore sought to evaluate the effect of BTZ treatment on the phosphorylation status of these key kinases. To accomplish this, Fibs were treated ± BTZ for 30 min, after which cultures were replaced with fresh medium and stimulated with TGF-β or FGF-2 and harvested after 30 min (**Figure 8A**). BTZ pretreatment significantly abrogated both the ability of TGF-β to increase phosphorylation of P38 (**Figure 8B**, left) and of FGF-2 to increase that of AKT (**Figure 8B**, right). This effect could be explained by inhibition of kinase activation and/or enhancement of phosphatase-mediated dephosphorylation. There is precedent for BTZ increasing *de novo* expression and/or activity of phosphatases (25, 32). To evaluate the importance of gene induction, we pretreated Fibs with 5,6-dichloro-1-β-D-ribofuranosylbenzimidazole (DRB), a reversible inhibitor of transcription, and then removed it and changed the medium prior to assessing the ability of BTZ to inhibit MyoFib differentiation (**Figure 8C**). Indeed, the ability of BTZ to attenuate TGF-β-induced α-SMA expression was significantly abrogated by pretreatment with DRB, implying a requirement for new transcription (**Figure 8D**).

**Figure 8.**
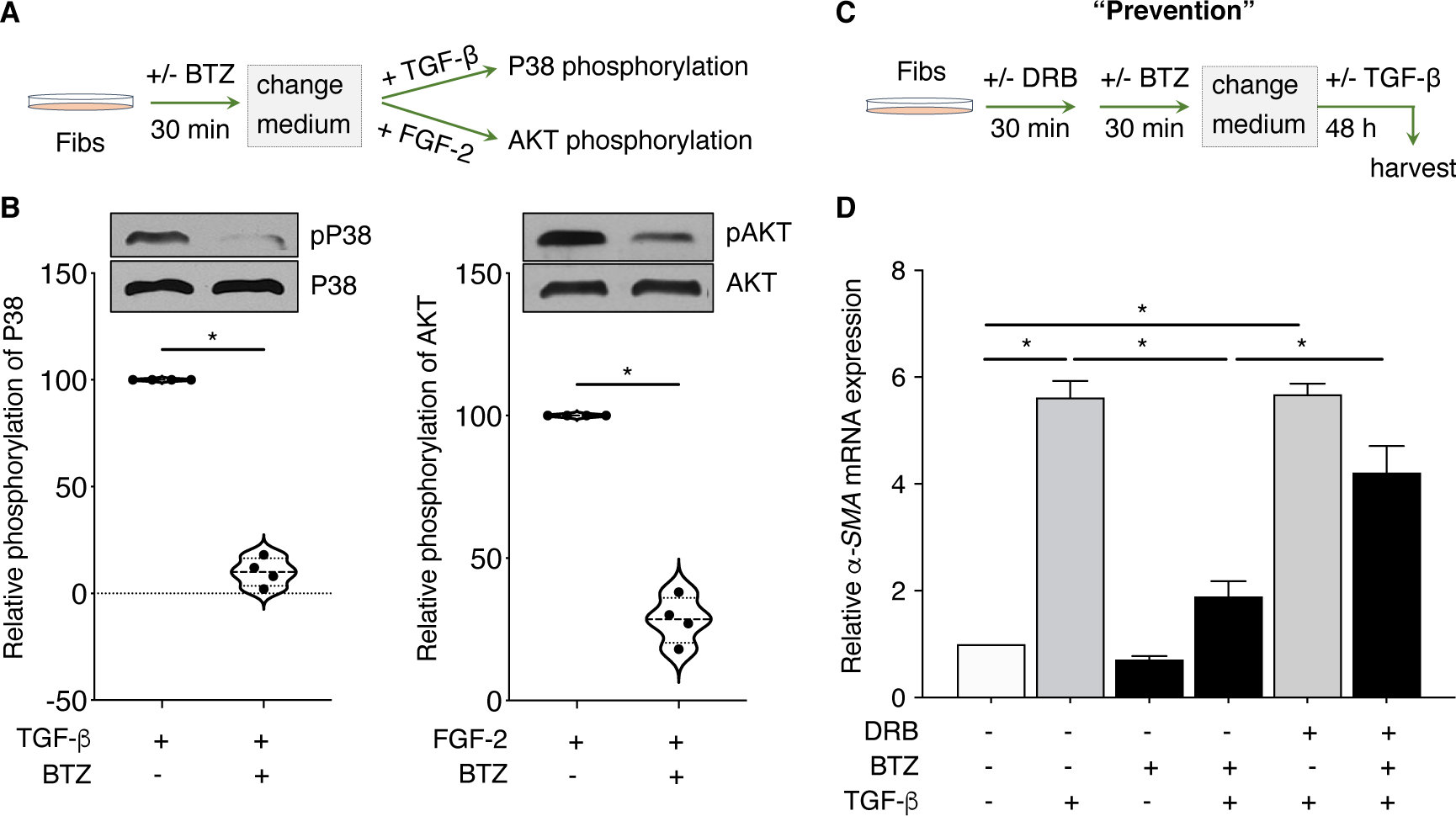
BTZ inhibits key kinases activated by TGF-β and FGF-2. (**A**) Design of experiments to assess BTZ effects on phosphorylation of key kinases activated by TGF-*β* or FGF-2. (**B**) Fibs were treated +/- BTZ (10 nM) for 30 min, after which the medium was replaced, and cells stimulated with TGF-*β* or FGF-2 for 30 min. Cells were harvested and phosphorylation status of P38 by TGF-*β* and AKT by FGF-2 was assessed by Western blot. Total P38 and AKT proteins were used to normalize phospho-P38 and phospho-AKT, respectively. Graphs represent mean densitometric analysis of phospho-P38 and phospho-AKT Western blots. (**C**) Design of experiments to assess ability of DRB to prevent BTZ modulation of TGF-*β*-induced MyoFib differentiation. (**D**) Cells were treated +/- DRB (25 μM) for 30 min and then treated +/- BTZ (10 nM) for an additional 30 min. Medium was replaced, and cells stimulated with TGF-*β* to analyze the expression of *α-SMA* mRNA by qPCR at 48 h. All data represent mean values (± S.E.) from 4 independent experiments. *P < 0.05, two-way ANOVA.

### Induction of dual-specificity phosphatase DUSP1 is uniquely associated with the inhibitory actions of BTZ on MyoFib differentiation

A publicly available RNA-seq database for human lung Fibs (33) identified a total of 90 phosphatase genes in these cells. We designed specific primer sets for each of these (see **Supplementary Table 2**). We treated Fibs with BTZ for 30 min and harvested them for phosphatase gene expression studies (**Figure 9A**, top). Of the 90 phosphatases screened by qRT-PCR, dual-specificity phosphatase *DUSP1* was the only one induced by BTZ, at a level of ∼2-fold (**Figure 9A**, bottom). The induction of *DUSP1* mRNA was rapid and was paralleled by that of its protein level (**Figure 9B**). Induction of functional DUSP1 protein was thus rapid enough to account for the rapid dephosphorylation depicted in **Figure 8A**. As was true for its ability to abrogate the BTZ inhibition of MyoFib differentiation (**Figure 8D**), the transcription inhibitor DRB likewise abolished the induction by BTZ of DUSP1 itself (**Figure 9C**). Together, these data suggest that DUSP1 is unique among Fib protein phosphatases in being transcriptionally induced by BTZ, and new gene induction was integral to the ability of BTZ to inhibit MyoFib activation.

**Figure 9.**
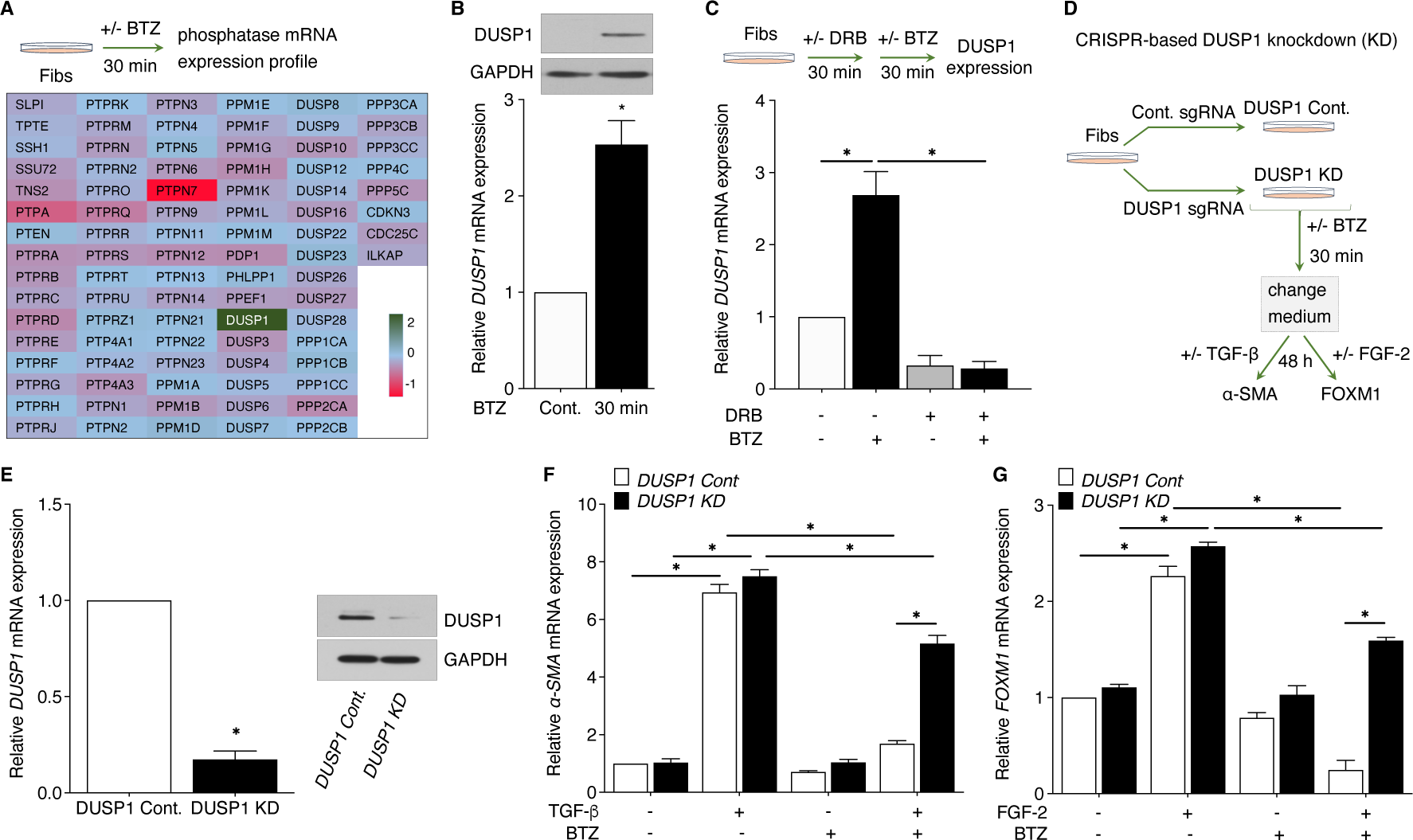
The anti-fibrotic actions of BTZ require *de novo* induction of DUSP1. (**A**) (Top panel) Design of experiment to assess the expression of phosphatase genes by BTZ. (Bottom panel) Fibs were treated +/- BTZ (10 nM) for 30 min, and cells were harvested immediately to assess the expression of various cellular phosphatase mRNAs by qPCR. (**B**) Cells were treated +/- BTZ (10 nM) for 30 min, harvested and assessed for the expression of *DUSP1* mRNA by qPCR (below) and protein by Western blot (above). (**C**) (Top panel) Design of experiment to assess the requirement for new transcription of the induction of DUSP1 by BTZ. (Bottom panel) Cells were treated +/- DRB (25 μM) for 30 min and then +/- BTZ (10 nM) for additional 30 min; cells were harvested immediately to assess the expression of *DUSP1* mRNA by qPCR. (**D**) Experimental design illustrating sgRNA-based *DUSP1* KD in MRC5 Fibs and assessment of BTZ capacity to attenuate actions of TGF-*β* and FGF-2. (**E**) Efficiency of KD of *DUSP1* by mRNA (left) and protein (right) determined by qPCR and Western blot, respectively, at 48 h post sgRNA transfection. (**F-G**) *DUSP1* Cont and *DUSP1* KD cells were treated +/- BTZ (10 nM) for 30 min, after which the medium was replaced, and cells stimulated +/- TGF-*β* (**F**) or +/- FGF-2 (**G**) and harvested at 48 h to analyze the expression of *α-SMA* or FOXM1 mRNA respectively by qPCR. All data represent mean values (± S.E.) from 3 independent experiments. *P < 0.05, two-way ANOVA.

### DUSP1 is required for BTZ-induced suppression of Fib activation

To specifically assess the role of DUSP1 in BTZ inhibition of Fib activation, we utilized CRISP-Cas9 KD of DUSP1 using MRC5 lung Fibs (**Figure 9D**). Using DUSP1 sgRNA, we generated Fibs deficient in DUSP1. Fibs receiving control (or non-targeting) sgRNA were designated as DUSP1 Cont. Fib KD of DUSP1 was confirmed at mRNA and protein levels (**Figure 9E**). Next, we assessed the impact of DUSP1 KD on the capacity of BTZ to inhibit TGF-β- or FGF-2-induced Fib activation. TGF-β and FGF-2 increased the expression of α-SMA and FOXM1, respectively, in DUSP1 Cont Fibs, and these actions were opposed by BTZ treatment (**Figure 9F-G**). These inhibitory actions of BTZ were significantly abrogated in DUSP1 KD Fibs. Collectively, these findings suggest an important role for DUSP1 in BTZ inhibition of Fib activation.

## Discussion

Because tissue fibrosis is characterized by elevated proteasome activity (5, 6) and BTZ is a clinically available proteasome inhibitor (34), the impetus to explore the potential of this agent in animal models of fibrosis is evident (14, 15, 35, 36). However, data in the lung have been limited, with both its safety and efficacy coming into question. Moreover, the relationship between the anti-fibrotic actions of BTZ and proteasomal inhibition has not been explicitly interrogated in any tissue or in Fibs in general.

Mutlu, et. al. reported that BTZ abrogated lung fibrosis at day 21 when dosed at days 7 and 14 post-bleomycin (17). By contrast, Fineschi, et. al. reported a lack of anti-fibrotic action but diminished survival in mice dosed daily with BTZ for the first 15 days after administration of bleomycin at the high dose of 4 units/kg (16). It is possible that both toxicity as well as lack of efficacy in the latter study were the consequence of the early and frequent dosing of BTZ along with the high dose of bleomycin employed. Importantly, neither of these studies examined the effect of BTZ on lung proteasomal activity. Semren et al. (37) studied the effects of oprozomib, a proteasome inhibitor which is an analog of the FDA-approved agent carfilzomib, in bleomycin-induced pulmonary fibrosis. It too was associated with accelerated weight loss and death, while lacking anti-fibrotic efficacy. The authors of this study advised caution regarding the prospects for proteasome inhibitors in the treatment of pulmonary fibrosis. Our first goal, then, was to attempt to resolve the therapeutic uncertainty regarding BTZ. Our protocol of BTZ dosing every three days beginning at day 9 resulted in a marked diminution of all indices of bleomycin-induced lung fibrosis. In addition, no mortality or histologic evidence of inflammatory cell recruitment or altered lung architecture was attributable to BTZ itself.

Although it has been generally assumed that potential anti-fibrotic actions of BTZ in the lung and other organs *in vivo* are attributable to its proteasomal-inhibitory actions, this assumption has not actually been directly validated. We were therefore surprised to see that the robust abrogation of pulmonary fibrosis in our protocol was dissociated from any demonstrable proteasome inhibition, as determined from both direct measurements of chymotrypsin-like peptidase activity and of global protein ubiquitination in lung tissue. At the same time, however, we are aware of no coherent mechanism ever articulated by which proteasome inhibition was envisioned to explain the anti-fibrotic actions of this agent. It is relevant in this regard to note that PGE_2_ and its downstream signaling intermediates cyclic AMP and PKA, which have been amply demonstrated to exert anti-fibrotic effects *in vivo* and *in vitro* (5, 27, 29, 38), have actually been reported to increase proteasomal activity (39–43). Likewise, azithromycin was reported to exert anti-fibrotic actions in lung Fibs *in vitro* and in a bleomycin model *in vivo* while enhancing proteasome activity (44). Both of these reports lend credence to our findings of a dissociation between anti-fibrotic actions and proteasomal inhibition. Recognizing that proteasome-independent actions of BTZ are appreciated (45–48), these unexpected findings prompted us to explore potential anti-fibrotic actions of BTZ in Fibs and in MyoFibs *in vitro* under conditions in which it likewise did not inhibit the proteasome.

We settled on a regimen involving treatment with 10 nM BTZ for 30 min because it had negligible impact on cellular proteasome activity determined at time points ranging from 30 min-24 h later and was unassociated with cytotoxicity up to 72 h later. In prior *in vitro* studies, Mutlu et al (17) and Fineschi et al (16) treated lung Fibs with BTZ at 200 nM and 1 μM, respectively, and employed continuous exposure without a subsequent change of medium prior to cellular activation endpoint analysis at time points ranging from 24 h to 48 h. No previous studies in Fibs or MyoFibs have explored such transient exposures or such low doses as ours, and one wonders if the cytotoxic effects of long- term continuous BTZ at the doses employed by others may have contributed to its previously reported *in vitro* suppressive effects. Using this protocol for BTZ treatment, we carried out the most comprehensive analysis to date of its actions on a variety of activation phenomena in lung Fibs and MyoFibs.

We found robust inhibition of mitogen-induced Fib proliferation as well as of activation of the transcription factor FOXM1 and expression of FOXM1-dependent proliferation- associated genes. Likewise, BTZ prevented the capacity of TGF-β to increase expression of characteristic MyoFib genes α-SMA and Col1α2. While prevention of MyoFib differentiation might halt further progression of a fibrotic disorder, reversing established fibrosis may require clearance of the activated MyoFibs that have already accumulated. Although MyoFibs were historically considered terminally differentiated cells, it is now well-recognized that they can be phenotypically de-differentiated – defined by a loss of α-SMA – back to or towards undifferentiated Fibs (27, 49, 50). Such de-differentiation has the potential to restore their sensitivity to apoptosis, and it has recently been suggested that MyoFib de-differentiation may be necessary for the resolution of fibrosis (51). We investigated the ability of BTZ to promote de- differentiation in two types of MyoFibs –those generated *in vitro* by differentiation of normal lung Fibs with TGF-β and those derived from the lungs of patients with biopsy-proven IPF. High baseline expression of fibrotic markers α-SMA and COL1α2 in these IPF Fibs was confirmed as we reported previously (5, 29). BTZ treatment resulted in robust de-differentiation of both types of MyoFibs. To our knowledge, this is the first demonstration of the *in vitro* de-differentiation potential of BTZ. In our de-differentiation studies, we consistently observed near-complete loss of COL1*α*2 protein to an extent that exceeded the reduction in either *COL1α2* mRNA or α-SMA protein. Under most circumstances, a proteasome inhibitor such as BTZ would be expected to prevent protein degradation. The underlying mechanism for this disproportionate loss of COL1*α*2 protein will require additional investigation in future studies.

It was imperative to determine if the de-differentiation of MyoFibs achieved by BTZ treatment did indeed sensitize cells to apoptosis. Apoptosis resistance in MyoFibs has been associated with an imbalance favoring expression of survival relative to apoptosis genes (52, 53). As expected, Fibs but not TGF-β-generated MyoFibs exhibited sensitivity to apoptosis elicited by activation of the classic death receptor FAS. However, de-differentiation of MyoFibs with BTZ sensitized them to a level of FAS-mediated cell death that was comparable to that exhibited by Fibs. This apoptosis sensitization was associated with decreased expression of the FOXM1-dependent survival gene *BIRC5*, but increased expression of the pro-apoptotic genes *APAF1* and *BID* and of *FAS* itself. BTZ sensitization of MyoFibs to FAS-mediated apoptosis was also recapitulated in IPF MyoFibs, which are notoriously resistant to apoptosis. Thus, our data demonstrate that de-differentiation of MyoFibs by BTZ results in a reprogramming of genes influencing apoptosis in a manner that restores their sensitivity, a phenomenon that might facilitate fibrotic Fibs or MyoFib clearance from fibrotic lungs. It will be critical to explicitly evaluate this possibility *in vivo* in future studies.

Our *in vitro* and *in vivo* data demonstrating that the anti-fibrotic actions of BTZ were independent of proteasome inhibition compelled us to seek to identify alternative mechanisms for these actions. To our knowledge, the possibility of proteasome-independent anti-fibrotic actions of BTZ has not been previously considered or supported. We considered that BTZ might activate the PGE_2_-cAMP axis, which exerts broad anti-fibrotic actions (5, 27, 29, 38), but our experimental data did not support generation of PGE_2_ and/or its downstream second messenger cAMP as a consequence of BTZ treatment. We have previously provided evidence that the kinases AKT and P38 play critical signaling roles in the activation of Fibs. The finding that BTZ dephosphorylated and hence inactivated both of these kinases was therefore an attractive clue to its Fib-inhibitory actions. The rapidity with which BTZ caused kinase dephosphorylation suggested the activation of a phosphatase, and the dependence on new transcription suggested induction of one or more phosphatases. Although BTZ has been shown in other cell types to induce a number of phosphatases (25, 54) – including DUSP1 – remarkably, DUSP1 was the only one of 90 phosphatases known to be expressed in human lung Fibs to be induced at the mRNA level by BTZ. The finding that loss of DUSP1 prevented the inhibitory effects of BTZ on TGF-*β*-induced MyoFib differentiation and FGF-2-induced FOXM1 expression conclusively established that induction of DUSP1 was required for these actions of BTZ. DUSP1 is known to dephosphorylate and thus inhibit activation of both P38 and AKT (55, 56). Although there is some precedent for DUSP1 mediating suppressive effects in mesenchymal cells and acting as a brake on tissue fibrosis (57), to our knowledge, this is the first such demonstration of this phenomenon in lung Fibs. Future work, including the generation of mice with a Fib-specific deletion of DUSP1, will be necessary to validate the role of this Fib phosphatase in limiting pulmonary fibrosis and to better understand its regulation and actions *in vivo*.

In this study, we report for the first time that BTZ abrogates experimental pulmonary fibrosis *in vivo* and diverse indices of lung Fib and MyoFib activation *in vitro* via a proteasome-independent mechanism. Such effects are instead attributable, at least in part, to rapid transcriptional induction of the phosphatase DUSP1, which opposes activation of the key pro-fibrotic kinases P38 and AKT. The ability of BTZ to promote MyoFib de-differentiation would be expected to concomitantly overcome the baseline resistance of these effector cells to apoptosis, which in turn represents a potential approach to reducing the accumulation of these pathogenic MyoFibs in the fibrotic lung. BTZ is the first-line treatment for multiple myeloma via subcutaneous dosing once or twice weekly and is relatively well tolerated. BTZ had promising beneficial effects in a small group of patients with pulmonary graft versus host disease following stem cell transplant, a condition characterized by fibrotic changes in the small airways and associated with TGF-*β* activation (58). A clinical trial of BTZ in scleroderma-associated pulmonary fibrosis is currently in progress (ClinicalTrials.gov: <u>NCT02370693</u>). Our results provide further support for and new mechanistic insights into the repurposing of BTZ for the treatment of IPF, and potentially other fibrotic disorders.

## Acknowledgments

We would like to thank Mikel Haggadone, Daniel Schneider, and Steven Huang for their valuable comments and suggestions that helped us to improve the quality of this manuscript.

## Competing financial interests

The authors declare no competing financial interests.

## Notes

This work was supported by National Institutes of Health grants HL94311 and HL144979 (to M. P.-G.), a Galvin Pilot Grant (to L.R.K.P.) and American Cancer Society Grant PF-17-143-01-TBG3 (to J.M.S).

### Competing Interest Statement

The authors have declared no competing interest.

