## Supplemental file for "Bortezomib inhibits lung fibrosis and fibroblast activation without proteasome inhibition"

**Supplementary Table 1.** Primer sequences used for qPCR

| Gene | Forward primer | Reverse primer |
|---|---|---|
| **Human sequences** | | |
| *GAPDH* | CAGCCTCAAGATCATCAGCA | ACAGTCTTCTGGGTGGCAGT |
| *FOXM1* | GACATCTATACGTGGATTGA | GGTGAATGGTCCAGAAGGAG |
| *CYCB1* | TGTGGATGCAGAAGATGGAG | GTGACTTCCCGACCCAGTAG |
| *CYCD1* | CGTGGCCTCTAAGATGAAGG | CCACTTGAGCTTGTTCACCA |
| *PLK1* | AGTCGACCACCTCACCTGTC | CTGACCATTCCACCAAGGTT |
| *BIRC5* | CCACTGAGAACGAGCCAGAC | GACAGAAAGGAAAGCGCAAC |
| *α-SMA* | ATCACCAACTGGGACGACAT | CATACATGGCTGGGACATTG |
| *Col1α2* | GGGTGAAATTGGAGCTGTTG | ACCAGTAAGGCCGTTTGCTC |
| *APAF1* | GGAGTGCATTGGGTTTCAGT | GAGAGACCTTGGGTGTTTGC |
| *BID* | TGTGAACCAGGAGTGAGTCG | TCTCTGCGGAAGCTGTTGTC |
| *FAS* | CGGACCCAGAATACCAAGTG | TGGTGAGTGTGCATTCCTTG |
| **Mouse sequences** | | |
| *β-Actin* | GACGGCCAGGTCATCACTAT | GCACTGTGTTGGCATAGAGG |
| *Col1α1* | ACCTCAGGGTATTGCTGGAC | CACCACTTGATCCAGAAGGA |
| *Ctgf* | GCAGACTGGAGAAGCAGAGC | ACACTGGTGCAGCCAGAAAG |
| *Tgf-β1* | GGAGAGCCCTGGATACCAAC | ATCCACTTCCAACCCAGGTC |

**Supplementary Table 2.** Primer sequences used for qPCR analysis of phosphatase mRNAs

| Gene | Forward primer | Reverse primer |
|---|---|---|
| **Human sequences** | | |
| *SLPI* | CAGCCTCAAGATCATCAGCA | ACAGTCTTCTGGGTGGCAGT |
| *TPTE* | ATGTCACTCTCCTCCTTGCC | ACTGCTGTCTCCCTTCTACA |
| *SSH1* | CGGTACATGGTGGTGGTGTA | CTCCACAGTCGGAGAACCAT |
| *SSU72* | AGCTCTGCCAGTGTATCCAG | TCAGTAGAAGCAGACGGTGT |
| *TNS2* | GCACTGGCCACTCTTACCAT | GAGAGGGCTGCTGTTCATTC |
| *PTPA* | GCTGCTTTCCTCTGCTGTCT | TCTGGAGTTTCCGCATAACC |
| *PTEN* | CACGACGGGAAGACAAGTTC | GGTTTCCTCTGGTCCTGGTA |
| *PTPRA* | GAAGGCCTGTAACCCTCAGT | CAAAGCCATACACGTCCACC |
| *PTPRB* | ATCCCTGGCAACAACTTCAG | TCAACACACTGGGTCACCAT |
| *PTPRC* | ACCTGGAATCCCCCTCAAAG | GACACATTGCAGCACTTCCA |
| *PTPRD* | CACACCACTGCATTCTCACC | GAAGGTGAGGCGTCTGTACT |
| *PTPRE* | TCAACCGAGTGATCCTTTCC | CCAGAAGTCCTCAACCGTGT |
| *PTPRF* | GTCCTGGAGCTCAGCAATGT | GGTGGCAGTTGTCTCTGTCA |
| *PTPRG* | AGGGACAATTCTGCACTGGA | CGATAATAGCTGCCCAGGGA |
| *PTPRH* | TGTCAGCATCTCCACAGTCC | CCATGAGACCCAGTAGCTGT |
| *PTPRJ* | ACATCACAGGCTTACGTCCA | GGATCGGCTCTGTGATGACT |
| *PTPRK* | TTGCCGCTTCCTTCAGATTG | ATGTAGGCCCAACACCAAGA |
| *PTPRM* | AGGGTGAACATGGTGCAAAC | CTAACTTGGGAAGCAGGCAC |
| *PTPRN* | CTCAGGCAGCTTCATCAACA | GATTTGGAGCCCTGTCTGTG |
| *PTPRN2* | GGCAATGGACTTTTACCGCT | CAGGTAGGTTTTCGGGAGGT |
| *PTPRO* | CGCCTTGTCAAGCTAGAACC | TGGAGCCAACTTGTTGACAG |
| *PTPRQ* | CCATCAGGTCGCATTTTGGA | TGGAACATCTGGTGGAGTGA |
| *PTPRR* | GTGGCTGCAGCTTTAGGACT | GAACTTCCTCAGAGGGCAGA |
| *PTPRS* | TCAGCGCTTTGAGACGATTG | TGGACTGTGATCTCCCCAAC |
| *PTPRT* | CACACCACTGCATTCTCACC | GAAGGTGAGGCGTCTGTACT |
| *PTPRU* | AGGCCCAGTACGATGACTTC | CATGCTGGGAAGTGTTGACC |
| *PTPRZ1* | GTCAGCGGAGGAGTTTCAGA | GGTCCGCATCAAAGCAGTAG |
| *PTP4A1* | ACACACAATCCAACCAATGC | GGCCAATCAAGAACATGGAT |
| *PTP4A2* | CTGTGTTGCAGTGCATTGTG | ATTGAACGCTCCCCTTCTTT |
| *PTP4A3* | GAAGGCCAAGTTCTGTGAGG | TCATCCCGCTCTCAATAAGG |
| *PTPN1* | CCAGTGGGTGAAGGAAGAGA | CCACGACCCGACTTCTAACT |
| *PTPN2* | AGAGTGGCCAAGTTTCCAGA | TGGCAGCATGTGTTAGGAAG |

| | | |
|---|---|---|
| *PTPN3* | ACACCACCCGGGTATTATTG | GACAACCTGCCAAAACTGTG |
| *PTPN4* | TGCTGGCAGAACCTACAATG | CAACAAGACTTGCCCCTGAT |
| *PTPN5* | CCACAAACCTCGTCTCCTCT | GACAGAAAGACCAGCAGCAG |
| *PTPN6* | CAAGTACCCGCTGAACTGCT | CAGAAAGCACGAAGTCTCCA |
| *PTPN7* | CCATCTTGCCAAATCCCCAG | CCACACCATCTCCCAGAAGT |
| *PTPN9* | GTGATTGTCATGACCACCCG | TCTCCACGCCTAGATTGGTC |
| *PTPN11* | CAAAGGGGAGAGCAATGACG | GTGTTAAGGGGCTGCTTGAG |
| *PTPN12* | ATTCACCTCCTCCCCTACCT | TAGTGAATACCTCCCGCTGG |
| *PTPN13* | AGGTTTGAATCCAGCAGTGG | CTTCCTCTTGCCGTTTTAGC |
| *PTPN14* | GCCTGTGAAGGAGAGACCTG | GGTGGCATCAACTCGATTCT |
| *PTPN21* | AGGAAGTGGCTACCAGAGCA | GTTGAGCACTCCCCATCAAC |
| *PTPN22* | GATTGTATGCAGGCCCAATC | TGGTGGGCAAGAATTACAGA |
| *PTPN23* | TCTGCTATGAGGCAGTGGTG | AAGGTGGTTCTTCTGGCTGA |
| *PPM1A* | GAAGGAGGCAGAGTTGGACA | CATTCCTCTTGCTTGCCAAT |
| *PPM1B* | GTCTTGCTGGCAAGCGTAAT | TCATTTGCCTGAGAGCTTCC |
| *PPM1D* | GGGTGGTTCTTGGAATTCAG | CGATTCACCCCAGACTTGTT |
| *PPM1E* | GGAAGTCTGTCGGTTTCCAG | TCAGGGTTCACGGTGTCATA |
| *PPM1F* | CTCACATGGACTGCTGGAGA | ACAGGCAAGCAGCAGGTAGT |
| *PPM1G* | AGCTGCCTCGAGTTGCTAAG | TCTGCCTCCTCACTGCTGTA |
| *PPM1H* | ACAGGCTGGGCATACAAAAC | TCCCCTGGTCACTCCAATAG |
| *PPM1K* | TTGATGAGCCAATTCTGCTG | CTTTCCGTTTGCCAATCTGT |
| *PPM1L* | CCTATGATGAAGCAGGCACA | AGCGTTCCCATCTTTGTCAC |
| *PPM1M* | GCTGGAGCTACAGGAGGATG | CTGTGTGGGTCCTCTTGGTT |
| *PDP1* | ACTGATGGGTTGTGGGAGAC | TGCATCTGTCCCAGAGTCAC |
| *PHLPP1* | CTATGTGGGACCTGCCTGAT | GGGGTCCAGAGGAGCTAAAT |
| *PPEF1* | GGGCTGCTATTTTGGACCAG | CCTCGATTGCTGCCTTCTTC |
| *DUSP1* | TGCCTTGATCAACGTCTCAG | ACCCTTCCTCCAGCATTCTT |
| *DUSP3* | GCTTTGGCTCAAAAGAATGG | ATCTCACGGTTCTGCCTCAC |
| *DUSP4* | CATCACGGCTCTGTTGAATG | TCACGGCATCGATGTACTCT |
| *DUSP5* | ATGGATCCCTGTGGAAGACA | GGTAAGCCATGCAGATGGTG |
| *DUSP6* | CACTGGAGCCAAAACCTGTC | CAGTGACTGAGCGGCTAATG |
| *DUSP7* | AGGCCATCAGCTTCATTGAC | TAGGCGTCGTTGAGTGACAG |
| *DUSP8* | GCCTGACTTCATCTGCGAGA | ATGACTTGGCAGCTGGAGAG |
| *DUSP9* | CACTGGAGCCAGAACCTGTC | ACAGTGACGGTGACAGAACG |
| *DUSP10* | CCACTGACAGCAACAAGCAG | CAAGTAAGCGATGACGATGG |
| *DUSP12* | AGGCAATGGGATACGAAGTG | ACGGTAGTTGGGTCAACAGC |
| *DUSP14* | GATCCACAGTGTGAGCAGGA | GGCTTTCACCCAGTTGTACG |

| | | |
|---|---|---|
| *DUSP16* | CAGCGAGATGTCCTCAACAA | GCTGTCATTCACAGGCACAC |
| *DUSP22* | TCACTGACTTTGGCTGGGAG | ACTGCCGATACTGATGGACC |
| *DUSP23* | TGTTACCTGGTGAAGGAGCG | TCGTTCGCTGGTAGAACTGG |
| *DUSP26* | CAGGACATGGCTAACAACCG | TGGATGCTCATGTCAAAGGC |
| *DUSP27* | TCTGGCCCAAGCTCTACATT | TAGTAGTCGGGCCCAGTGTC |
| *DUSP28* | GCCTGCCTAGTCTACTGCAA | CGCTCTTCACCATCTGGAA |
| *PPP1CA* | CTCAAGATCTGCGGTGACAT | CTTGCCCCTGTCCACATAGT |
| *PPP1CB* | TGGTGGAATGATGAGTGTGG | CTGTTCGAGGTGGAGTGACA |
| *PPP1CC* | GCCATCGTGGATGAGAAGAT | GTCATTTTCACCCCAGCCTA |
| *PPP2CA* | AGGTGTTCACCAAGGAGCTG | CCTCTTGCACGTTGGATTCT |
| *PPP2CB* | GGTGTCATGATCGGAATGTG | GCTGGGTCAAATTGAAGGAA |
| *PPP3CA* | TGGCCAGAGTGTTCTCAGTG | TCAACAGTAGCGCTTTGCAG |
| *PPP3CB* | CAACCATGAATGCAGACACC | AGTGCAGCAAGAGGCAAACT |
| *PPP3CC* | GATCAGAGCCATTGGGAAGA | AGTCTGCTTGCCTCCTGAGA |
| *PPP4C* | CACGCAGGTCTATGGCTTCT | ACGCAGAAGATCTTGCCATC |
| *PPP5C* | GGAGCTCATGCAGTGGTACA | TTGAGTGTGGTTTCCACGAG |
| *CDKN3* | CATCATCATCCAATCGCAGA | CTCCCAAGTCCTCCATAGCA |
| *CDC25C* | TGTCAACCCAGAAACAGTGG | CTGGATGTGTCCTCCCAGAT |
| *ILKAP* | GTGTTTTGGGCGTGCTAGAG | CAGGCCAACAAAATGAACCT |

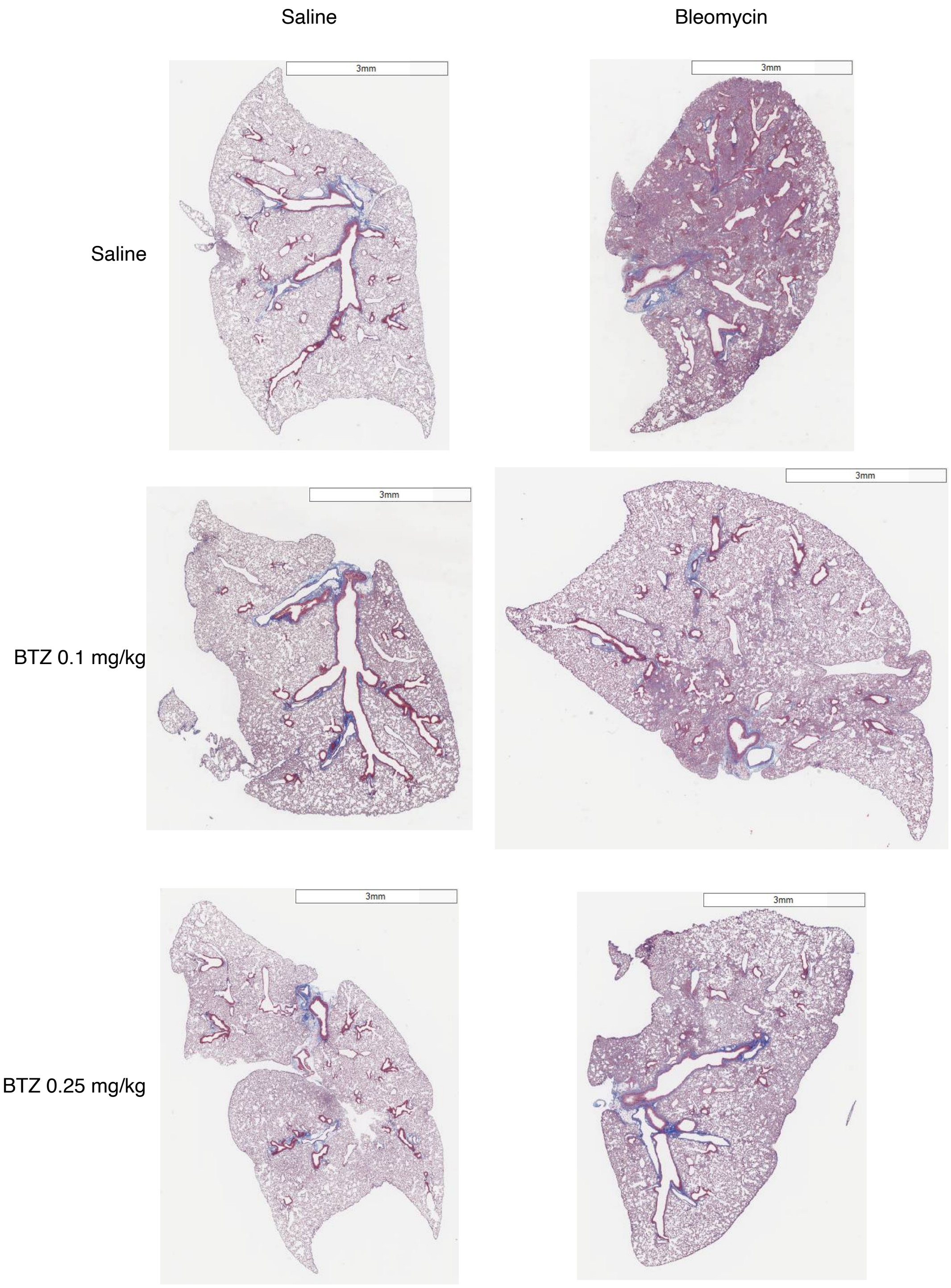


**Supplementary Figure 1. BTZ administration attenuates interstitial collagen in a mouse model of bleomycin-induced lung fibrosis.** Low magnification images of

Masson's trichrome-stained lung sections from mice administered saline or bleomycin treated and subsequently treated with or without BTZ. Scale bars: 3 mm.

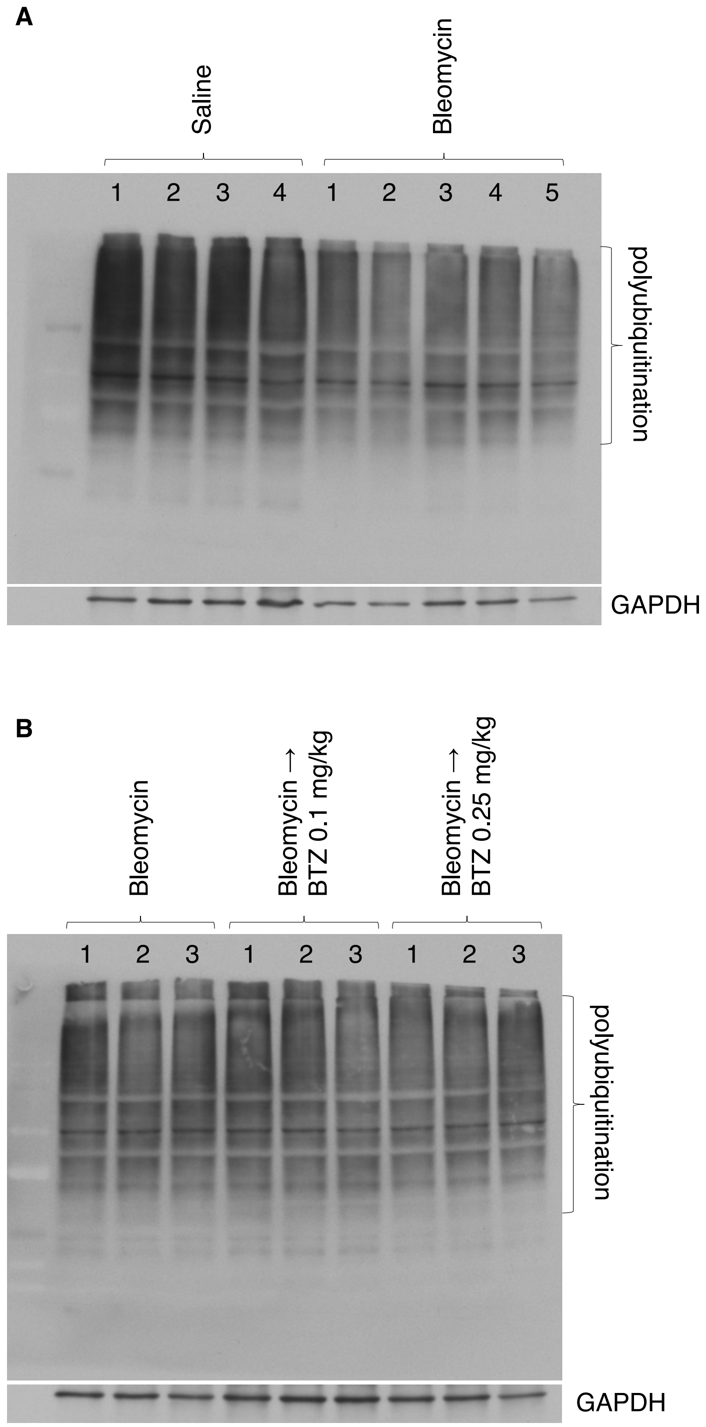


**Supplementary Figure 2. Global protein ubiquitination in lungs of mice treated with or without bleomycin and BTZ.** (**A**) Lungs were harvested on day 21 from mice treated on day 0 with bleomycin or saline; tissues were lysed, and global protein ubiquitination was determined by Western blot analysis. (**B**) Lungs were harvested on day 21 post-bleomycin from mice treated with or without BTZ at 0.1 mg/kg or 0.25 mg/kg; tissues were lysed, and protein ubiquitination was determined by Western blot analysis. Protein quantification was performed using ImageJ software and GAPDH protein was used to normalize ubiquitinated protein levels.

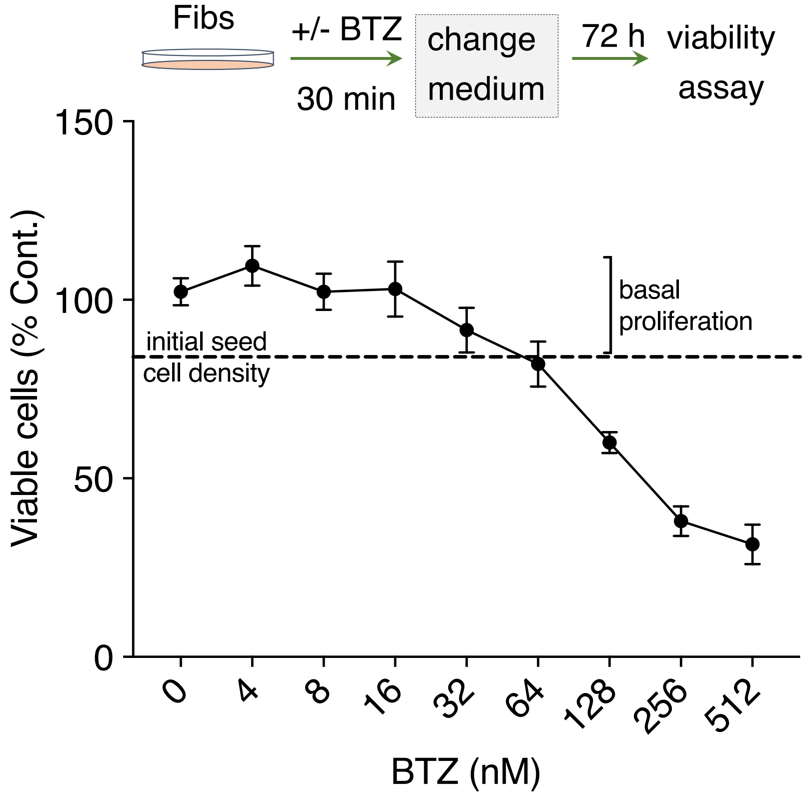


**Supplementary Figure 3. BTZ reduces Fib survival in a dose-dependent manner.**

(Top panel) A wire diagram to illustrate experimental design to assess Fib cell viability. (Bottom panel) Fibs were treated with increasing concentrations of BTZ for 30 min, after which the medium was replaced and cell viability was assessed at 72 h using the CellTiter-Glo luminescent assay kit (Promega).

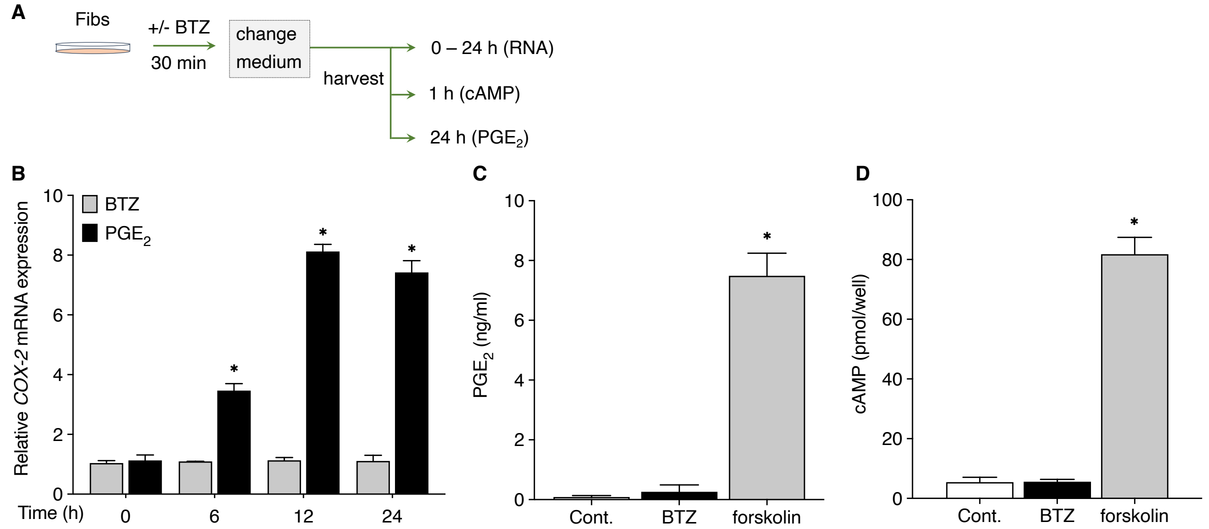


**Supplementary Figure 4. Inhibition of Fib activation by BTZ is independent of $PGE_2$ production or signaling.**

(**A**) A wire diagram to illustrate experimental design to assess BTZ capacity to modulate $PGE_2$ or its downstream second messenger, cAMP. (**B**) CCL-210 cells were treated +/- BTZ (10 nM) or $PGE_2$ (500 nM) for 30 min, after which the medium was replaced. Cells were harvested at 0, 6, 12 and 24 h time points, and assessed for the expression of COX2 mRNA by qPCR. (**C**) Cells were treated +/- BTZ (10 nM) or forskolin (500 nM) for 30 min, after which the medium was replaced. Conditioned medium was harvested 24 h later and assessed for levels of $PGE_2$ by ELISA. (**D**) Cells were treated +/- BTZ (10 nM) or forskolin (500 nM) for 30 min, after which the medium was replaced. Cells were harvested at 4 h and lysates assessed for intracellular levels of cAMP by ELISA. All data represent mean values (± S.E.) from 3 independent experiments. *P < 0.05, 2-way ANOVA.
